# Fabrication of mRNA encapsulated Lipid nanoparticles using *State-of-the-art* SMART-MaGIC Technology and transfection *in vitro*

**DOI:** 10.1101/2024.05.17.594777

**Authors:** Niloofar Heshmati, Jaidev Chakka, Yu Zhang, Mohammed Maniruzzaman

**Author notes:** Both authors contributed equally (Niloofar Heshmati, Jaidev Chakka). Corresponding Author Mohammed Maniruzzaman, Ph.D., Professor and Chair, Department of Pharmaceutics and Drug Delivery, School of Pharmacy, University of Mississippi, Oxford, MS 38655.

## Abstract

The messenger ribose nucleic acid (mRNA) *in lieu* of Corona virus of 2019 (COVID-19) vaccines were effectively delivered through Lipid nanoparticles (LNP) which were proved effective carriers for clinical applications. In the present work, mRNA (erythropoietin (EPO)) encapsulated LNPs were prepared using a next generation state-of-the-art patented Sprayed Multi Absorbed-droplet Reposing Technology (SMART) coupled with Multi-channeled and Guided Inner-Controlling printheads (MaGIC) technologies. The LNP-mRNA were synthesized at different N/P ratios and the particles were characterized for particle size & zeta potential (Zetasizer), encapsulation or complexation (gel retardation assay) and transfection (Fluorescence microscopy) in MG63 sarcoma cells *in vitro*. The results showed a narrow distribution of mRNA-lipid particles of 200 nm when fabricated with SMART alone and then the size was reduced to approximately 50 nm with the combination of SMART-MaGIC technologies. The gel retardation assay showed that the N/P>1 showed strong encapsulation of mRNA with lipid. The *in vitro* results showed the toxicity profile of the lipids where N/P ratio of 5 is the optimized with >50% cell viability. The functional LNP-mRNA were prepared and analyzed with SMART-MaGIC technologies which could be a potential new fabrication method of mRNA loaded LNPs.

**Highlights:**

1. The particle size was reduced to around 50 nm with implementation of SMART-MaGIC.
2. The loading efficiency is 100%.
3. The functionality of the mRNA is unaffected during the preparation process.
4. The transfection facilitated transient expression of the protein *in vitro*.
5. The EPO mRNA is more effective than EPO protein to reduce chemo-toxic effects *in vitro*.

## 1. Introduction

Messenger RNA is the intermediary molecule that transcribes DNA information and participates in the translation of protein in cytoplasm of the cell. The successful administration of mRNA based COVID vaccines has proved that mRNA could be a viable candidate with the potential to save millions of lives around the world^1,2^. Erythropoietin (EPO) is a glycoprotein cytokine that acts as hematopoietic growth factor (HGF) and stimulates erythropoiesis. The mRNA technology has been widely explored beyond vaccines to treat infectious and chronic diseases^3^. The recent demand for this technology raises new problems such as carrier compatibility, toxicity, and therapeutic efficacy of the delivery target. The immunogenicity and its susceptibility to RNases in the cell are the primary barriers for its application. These barriers are overcome by encapsulating the RNA molecules into lipid nanoparticles (LNPs) consisting of ionizing lipids, cationic lipids, PEG lipid and cholesterol as stabilizer^4–6^. The lipid composition with cationic and ionic lipids will drive the site specific delivery such as liver^7^.

The particle size plays a key role in nanoparticle-based therapeutics^8–10^. The fabrication process plays a crucial role to determine the particle size. The current techniques used to produce LNP carriers are microfluidic^11,12^ and jet mixing^13^-based processes for bulk manufacturing for commercial applications. Among the two, microfluidics is a very popular and widely used method for fabrication of particles. At the lab scale, solvent diffusion method^14^ or thin-film hydration^15^ methods are used for preparing the mRNA encapsulated LNPs. The particle size of the particles has a distinct impact in the immunogenicity of the formulations. The particle size of around 50 nm showed lower immunogenicity in mice^16^. The lower particle size will have better therapeutic potential thereby to escape reticular endothelial system in cancer treatment^17^.

The incorporation of innovative advanced technologies like three-dimensional printing (3D) over the last decade has presented distinctive opportunities for creating scaffolds used in biomedical applications^18–20^. This technological progress has made a notable contribution to the field of pharmaceutics and drug delivery by facilitating the design and production of tailored formulations for individual patients^21–23^. A multitude of investigations have delved into the potential applications of different categories of 3D printers in crafting various forms of dosage for diverse therapeutic compounds^24–27^. The introduction of particle formation through extrusion-based 3D printing offers several benefits^28^. It is adaptable to a variety of mechanisms for particle creation, thereby enhancing its adaptability. The coalescence of nanoparticles within a single production setup can be particularly advantageous for bioprinting, especially in scenarios involving the simultaneous delivery of multiple agents, such as growth factors^29^.

Our lab has invented a novel technology called Sprayed Multi-Adsorbed droplet Reposing Technology (SMART)^30,31^, which combines extrusion-based 3D printing with emulsion evaporation or solution dispersion techniques to produce nucleic acid-loaded lipid nanoparticles (LNPs)^32^. SMART overcomes limitations of conventional emulsion evaporation methods by utilizing shear force from a syringe nozzle instead of sonication energy, allowing for encapsulation of heat-sensitive agents and biomolecules without the need for emulsion or solution cooling. This platform enables rapid production of nanoparticles such as polylactide-co-glycolide and chitosan with precise dosage control, offering faster production compared to traditional methods. We developed plasmid DNA encapsulated lipid nanoparticles using SMART technology alone previously of around 150 nm^32^. Along with the SMART, we also developed another technology named Multi-channeled And Guided inner Controlling printhead (MaGIC, WO Patent-2024/026039) nozzles that has the ability to neutralize the pressures encountered by the particles and stabilize them to yield a small and uniform particle size with improved encapsulation efficiency.

In general, the MaGIC printheads are a series of custom-designed, multi-channel pneumatic printheads controlled by a linker, a designed functional system, and one or more outlets. To achieve uniform concentration mixing on the microscale, electro kinetics and surface tension are considered at small length scales, rather than inertial effects, resulting in turbulence and good mixing at macroscales^33^. In this study, a coaxially designed multi-channel MaGIC printhead was developed and applied for precise nuclei formation and controllable aggregation processes. While chip-based microreactors are known to benefit nanoparticle production, they often encounter issues with tube clogging due to the adhesion of precipitated particles to the tube walls^34,35,36^. Thus, the coaxially designed MaGIC printhead offers a solution to avoid clogging and facilitates easy continuous production. For further details on MaGIC printheads, two immiscible liquids were flowed through inner and outer tubes, maintaining an annular and laminar flow to create a micro-space between the outer fluid walls. The inner microchannel can be controlled using 3D printing technology, especially stereolithography printing. In this setup, nanoparticle reactions can be precisely controlled by the gradient concentration of reactants in the outer fluid, thereby preventing particle accumulation on the tube walls. MaGIC printheads are easy to design using CAD and 3D printing technology, offering a one-stop generation process with low cost and easy adjustment and maintenance.

In the present study, we encapsulated Erythropoietin (EPO) and green fluorescence protein (GFP) mRNAs in LNP with cationic lipids. The nanoparticle preparation, characterization, encapsulation efficiency and therapeutic efficacy *in vitro* were evaluated.

## 2. Experimental

### 2.1 Materials

The following lipids 1,2-distearoyl-sn-glycero-3-phosphocholine (18:0, DSPC), Cholesterol (Chol), 1,2-dimyristoyl-rac-glycero-3-methoxypolyethylene glycol-2000 (DMG-PEG) and 1,2-dioleoyl-3-trimethylammonium-propane (18:1, DOTAP) were procured from Avanti Polar, USA. All materials were used without any modifications. All materials were analytical grade and used without any modifications.

### 2.2 Messenger ribonucleic acid (mRNA)

The mRNA for green fluorescence protein (GFP, CleanCap EGFP mRNA) and Erythropoietin (EPO, CleanCap EPO mRNA) were purchased from Trilink BioTechnologies, USA. The mRNA sequences were provided in supplementary information (Sequence S1 and S2). The mRNA was prepared using CleanCap technology as per the manufacturer. The stock mRNA sequences were used as such without further changes.

### 2.3 Fabrication of Lipid-mRNA particles

The platform technologies SMART and MaGIC were employed for preparation of LNPs.

#### 2.3.1 SMART

SMART involves the extrusion of the lipid and mRNA components through a syringe fitted with 21 G nozzle (0.4 mm) at a pressure of 200 KPa into the MaGIC nozzle to ensure uniform particle size. Briefly, the lipids DSPC:Chol:DOTAP:DMG-PEG of 10:48:40:2 were mixed in ethanol solution at 6 mg/mL stock concentration. The mRNA was dissolved in 10 mM Citrate buffer with pH 6.0. The lipid solutions were taken into a 3 mL syringe and extruded at 200 KPa (BioX, Cellink, USA) into mRNA solution at 2:1 ratio to get a final lipid concentration of 2 mg/mL. The extruded mixture was taken back into a 3 mL syringe and extruded again to ensure complete encapsulation of mRNA. The final LNPs were dialyzed against 4x deionized water for 1 h using 3.5 KDa dialysis bag (Snakeskin Dialysis Tubing, ThermoFisher, USA). The particles were stored at 4 ^o^C for further studies. To ensure maximum ecapsulation or complexation the LNP and mRNA at different N/P ratios were analyzed.

#### 2.3.2 SMART-MaGIC

The particles fabricated with SMART and MaGIC technologies to optimize the particle size. The MaGIC nozzle was attached to the syringe loaded with LNP-mRNA and extruded through MaGIC nozzle. The optimized formulation using both SMART and MaGIC at different N/P ratios varying from 0.1 to 125 were prepared by varying the lipid quantities keeping the mRNA (1 μg/mL).

### 2.4 Characterization

The fabricated LNP-mRNA (EPO/GFP) nanoparticles were characterized for particle size and zeta potential using dynamic light scattering (DLS) method in Zetasizer (Malvern, USA). The stability of the particles was tested by gel retardation assay. A 2% agarose gel was prepared with 1 µg of EDTA. The LNP-mRNA particles were mixed with a 6x gel loading dye and a total 20 µL sample was loaded into each well for all the different N/P ratios. The gel was run at 80 v for 30 min. The gel was imaged under UV trans-illuminator and photographs were taken using camera. The images were processed for clarity using Microsoft PowerPoint (Microsoft, USA).

### 2.5 Encapsulation Efficiency

Encapsulation efficiency of mRNA within the LNPs was determined indirectly by quantifying unloaded mRNA (Quanti-iT RiboGreen Assay kit, Invitrogen, USA). The kit includes a dye that exclusively stains the free mRNA (non-complexed) in suspension. The following formula was used for the encapsulation efficiency (%): [(mRNA given-free mRNA)/mRNA given]*100.

### 2.6 Cell culture

To investigate the transfection efficiency and cell toxicity behavior of the formulations, the osteosarcoma cell line MG63 (ATCC, USA) was used. The cells were cultured in low glucose dubecco’s modified eagle medium (DMEM, Gibco, USA) supplemented with 10% fetal bovine serum (Gibco, USA) and 1% Penicillin-streptomycin (Gibco, USA). The cells were cultured in a CO_2_ incubator (CellExpert, Eppendorf, USA) at 37 ^o^C with 5% CO_2_. The cells were harvested using Trypsin-EDTA (Gibco, USA) method at 80% confluency. Cells were seeded in 96-well plates at a density of 10,000 cells/well (10^5^ cells/mL). Once the cells were attached, cell media was exchanged with fresh media containing LNPs at various concentrations to deliver the targeted dose of mRNA in each well. After 24 hours incubation, transfection efficiency as well as cells viability were investigated.

### 2.7 *In vitro* transfection

The LNP-mRNA (GFP) prepared using SMART-MaGIC were used for this study. Fluorescent microscopy was used to detect the expression of GFP. Cells were fixed prior to imaging using 3.7% formaldehyde for 20 minutes. The GFP fluorescence was imaged under fluorescence microscope equipped with multichannel laser system for detecting green wavelength (488 nm) (IX83, Olympus, USA). The fluorescent images were processed and merged with the brightfield images using ImageJ (v1.54g, ImageJ, NIH, USA).

### 2.8 Transfection Efficiency

The LNP-mRNA (EPO) prepared using SMART and MaGIC were used for this study. To monitor the expression of EPO, cell media was assayed using enzyme linked immunosorbent assay (ELISA, EPO, PicoKine, Boster, USA) as per the manufacturer’s instructions in duplicates.

### 2.9 Cell viability

The cell viability to reflect the toxicity effect of the formulations was evaluated in MG63 cells using MTT assay (MTT, Sigma Aldrich, USA). Briefly, after incubation, cell media was replaced with 0.5 mg/mL solution of MTT reagent in DPBS. After 2 hours incubation, formazan crystals formed by viable cells were dissolved in DMSO (Sigma Aldrich, USA) and their concentration was quantified by their absorbance at 540 and 700 nm using a multimode reader (Synergy H1, Biotek, USA).

### 2.10 Statistics

All experiments were carried out with triplicates. The data were shown as mean ± standard deviation. The significance of difference, *p<0.005 was evaluated doing One-way ANOVA (v5.03, GraphPad Prism, USA).

## 3. Results

### 3.1 SMART and MaGIC

The fabrication technology (Figure 1A, SMART and MaGIC) for LNPs was developed in-house. The technology was already tested and validated with our previous work (Jaidev and Maniruzzaman, Molecular Pharmaceutics). The MaGIC system (Figure 1B) was newly introduced in this work for the preparation of Lipid-mRNA nanoparticles.

**Figure 1.**
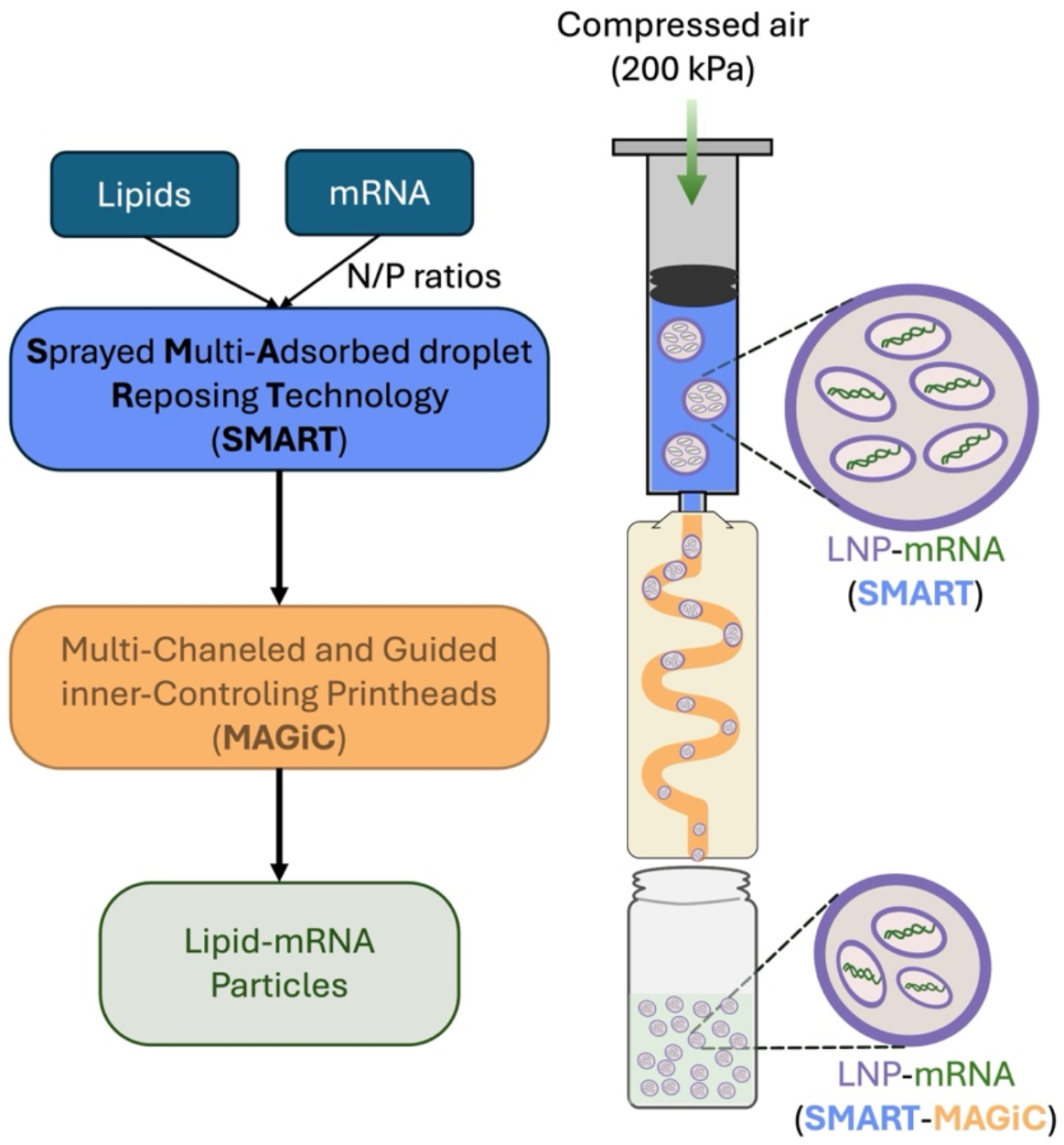
Schematic showing the process for formation of Lipid-mRNA nanoparticles using SMART-MaGIC technologies. The incorporation of MaGIC to the fabrication process has reduced the size of the LNP-mRNA particles.

### 3.2 Preparation of particles with SMART

Figure 1 showed the schematic of the SMART-MaGIC system that is used for the study. The SMART system is a 3D extrusion process and MaGIC is fabricated using the 3D printing process. The initial trials of the particle synthesis were carried out and the particle size and zeta potential were provided for the formulations at different N/P ratios from 0 to 100 (Figure 2). The extrusion process can be repeated to optimize the particle size. The results showed that there is no significance of difference between the particle size with one or two repeats (run1 and run2) of the extrusion process. The toxicity profile of the prepared formulations was studied in MG63 showed higher toxicity with respect to increased loading of particles within that N/P ratio (Figure 3). The N/P ratio 1 showed increased cell viability at all concentrations compared to N/P 10 and 100.

**Figure 2.**
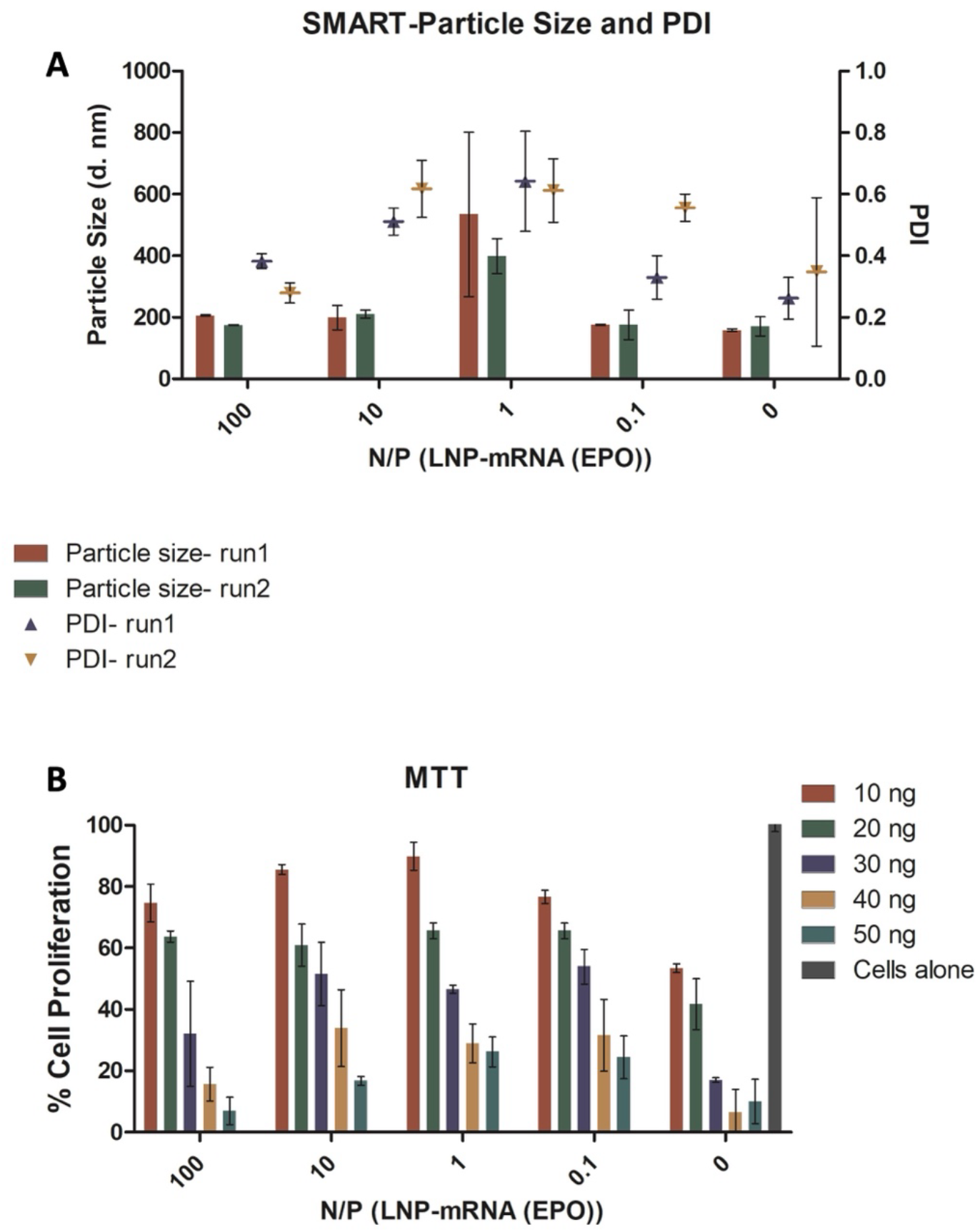
Fabrication of LNP-mRNA (GFP) using SMART system showing the [A] particle size and polydispersity index (PDI) and [B] *In vitro* cell viability of the formulations showing a dose-dependent cell viability. The lipid content of blank (0) is similar to N/P 100. All the data are represented as mean±standard deviation with n=3 replicates.

**Figure 3.**
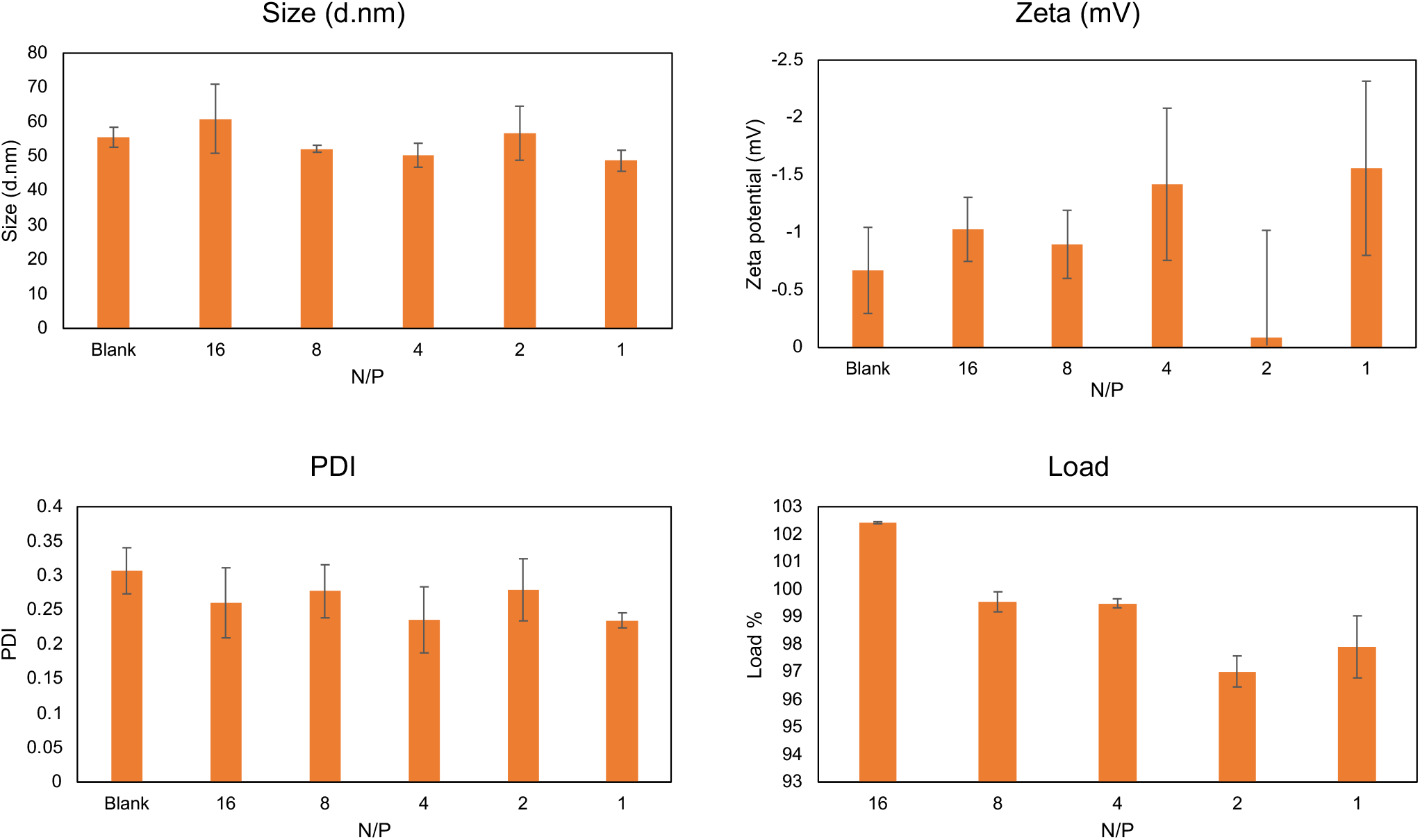
Characterization of LNP-mRNA (GFP) prepared using SMART-MaGIC system for their Particle size, Zeta potential, Polydispersity index, and Percentage loading of mRNA. All the data are represented as mean±standard deviation with n=3 replicates. The particle size for the least N/P 1 is around 50 nm is obtained using SMART-MaGIC process with around 100% loading of mRNA (EPO).

### 3.3 Preparation of particles with MaGIC

We narrowed the further exploration of the N/P ratio between 1 and 16 to optimize the formulation by preparing particles with SMART-MaGIC together. Figure 3 showed the particle size, polydispersity index, and zeta potential of the prepared formulations with different N/P ratios. For all the formulations, the particle size is around 50 nm and the zeta potential is in the negative range. The addition of lipid at higher N/P ratios showed comparative reduction in the negative charge. The addition of MaGIC has reduced the particle size for the formulations from more than 200 nm (Figure 1A) to around 50 nm. The technique showed a uniform and non-significant difference in the polydispersity index which could be the reason for a more consistent particle size across different N/P ratios. This could validate the method for the fabrication of different complexes of lipid nanoparticles and mRNA that will lead to a uniform particle size. The increase in mRNA loading was observed highest for N/P 16 and not much different between N/P 1 and N/P 4 & 8.

### 3.4 Comparison of formulations GFP vs EPO

The robustness of the particle preparation using SMART-MaGIC with different mRNA products (GFP and EPO) were studied by fixing the lipid content and varying the mRNA amounts as shown in figure 4. The N/P ratios are range between 0.625 and 10. The particle size is very much similar between GFP and EPO as shown in figure 4. The increased mRNA content showed increased complexation with mRNA providing a particle size closer to 80 nm. The N/P ratio of 5 was chosen to be the optimized formulation for which the zeta potential is around 4.59±0.6. Clearly the introduction of MaGIC along has also impacted the charge of the LNP-mRNA nanoparticles. The loading efficiency of mRNA with all the formulations is 100%.

**Figure 4.**
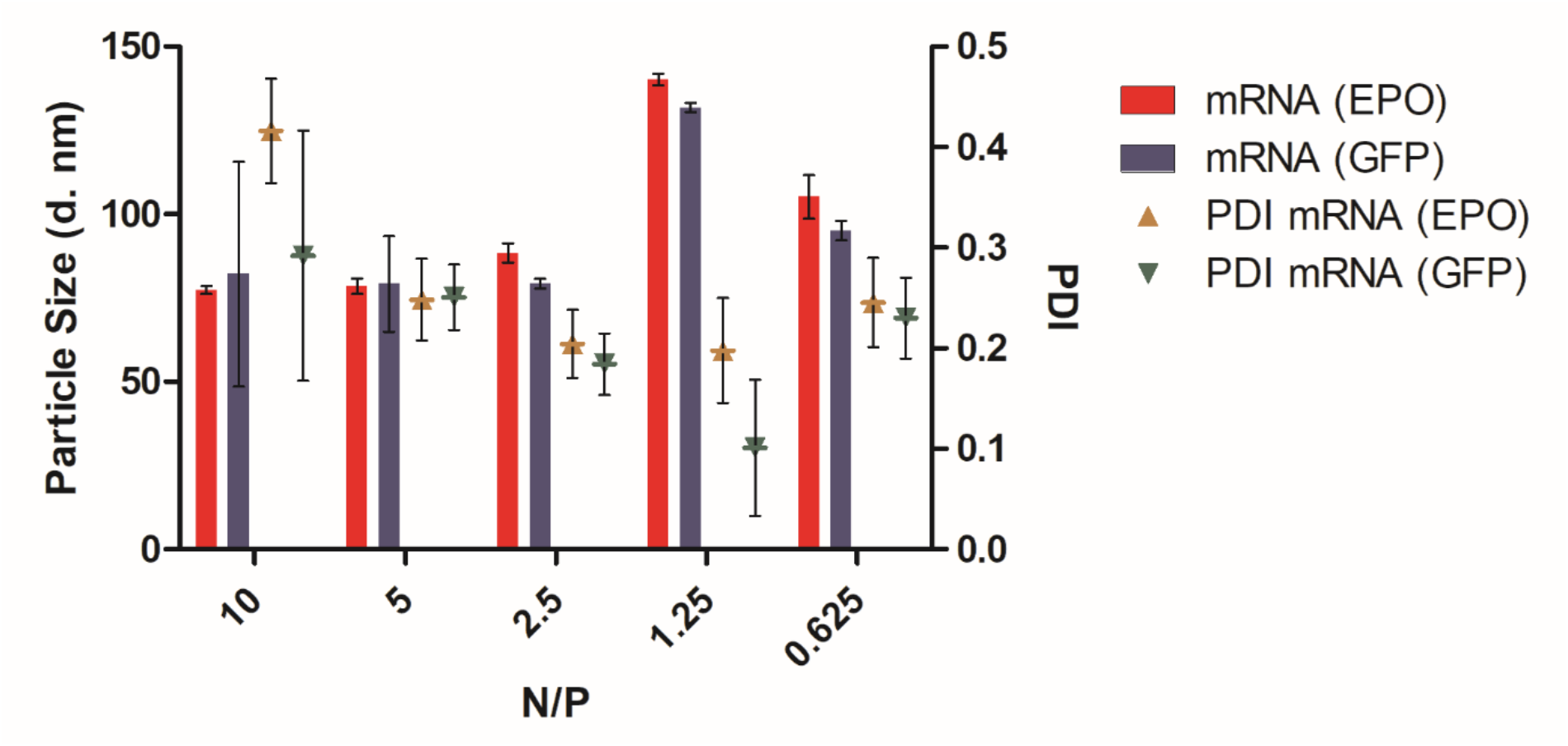
The image showing the particle size distribution of LNP-mRNA formulations at different N/P ratios encoding the GFP and EPO genes. The particle size was plotted on left axis and polydispersity index (PDI) was plotted on right axis. All the data are represented as mean±standard deviation with n=3 replicates.

**Figure 5.**
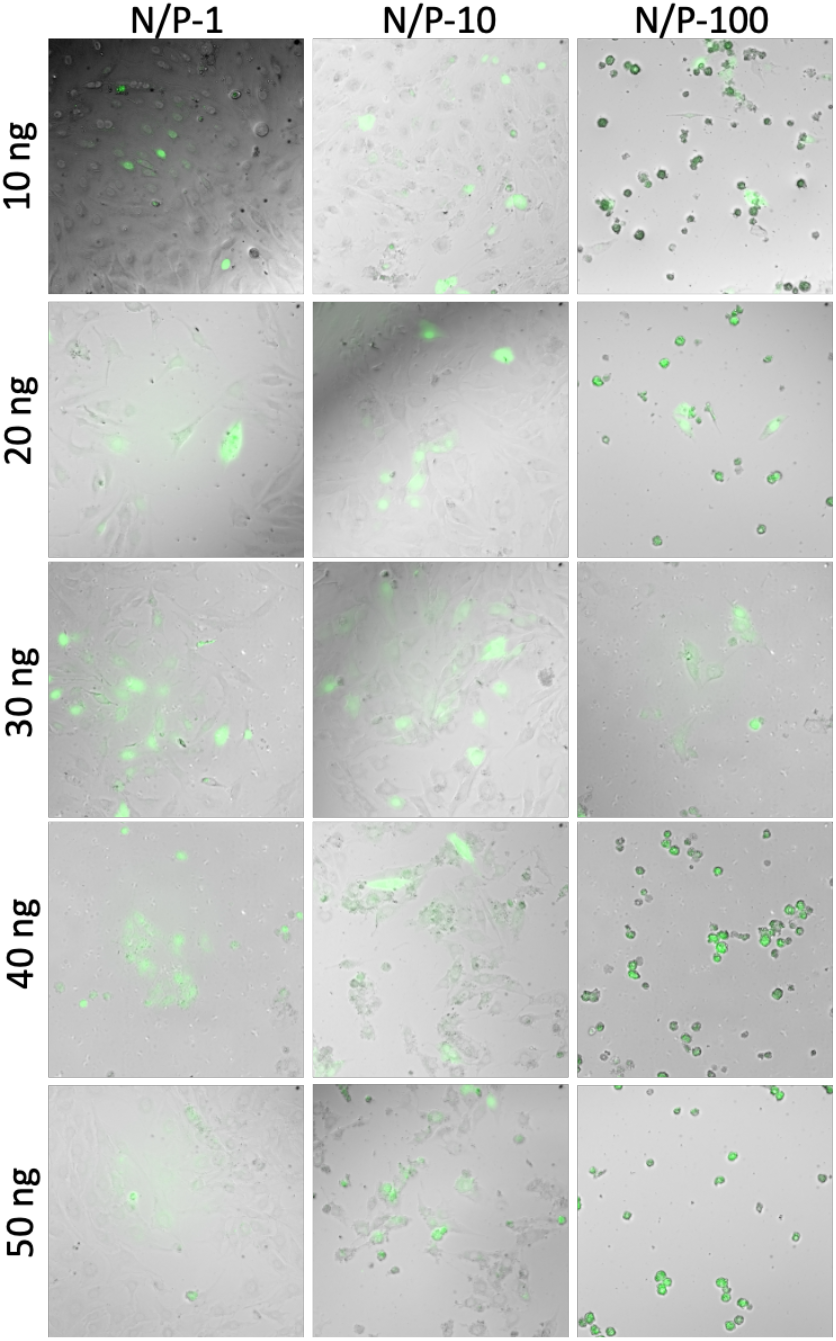
Transfection of LNP-mRNA (GFP) nanoparticles fabricated at different N/P ratios using SMART-MaGIC method at a dose dependent manner in MG63 osteosarcoma cells in a fluorescence microscope. The fluorescence (GFP) is overlayed on to the bright field image of the cells in ImageJ.

**Figure 6.**
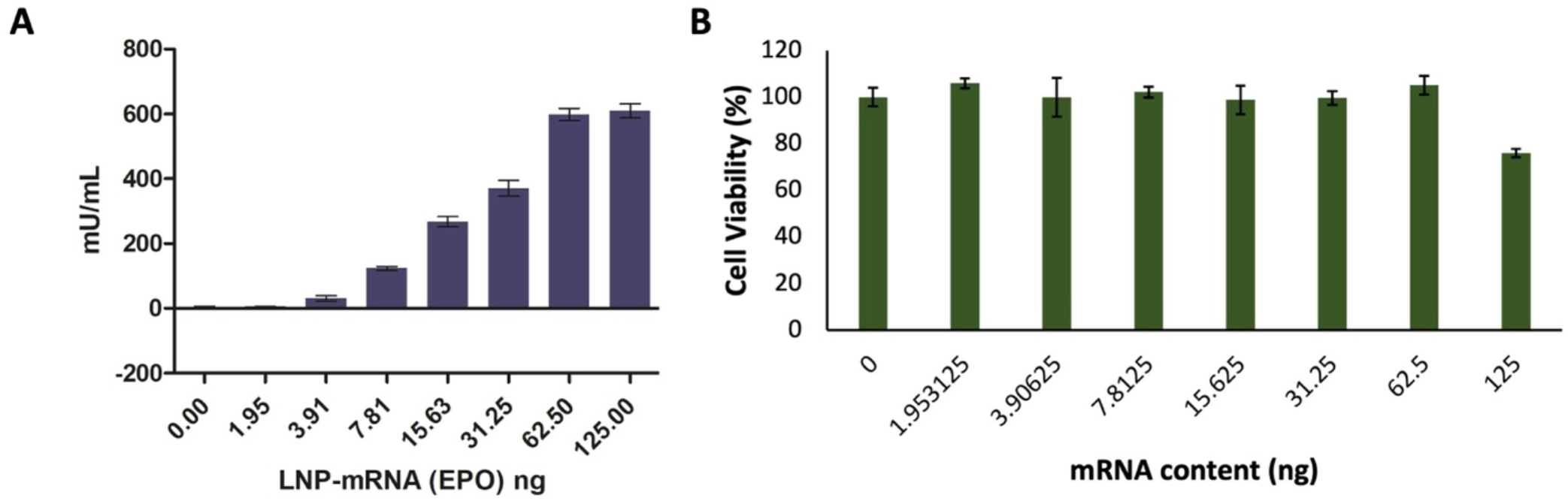
The image showing the [A] ELISA of LNP-mRNA formulations at N/P ratio 5 encoding the EPO gene and [B] Cell viability performed by MTT assay where the LNP-mRNA (EPO) dose was given based on the mRNA content. All the data are represented as mean±standard deviation with n=3 replicates.

### 3.5 *In Vitro* Transfection and EPO expression

The *in vitro* transfection in MG63 cells showed dose dependent transfection enhancement for N/P-1 and 10 while the N/P-100 showed toxicity with rounded cells in all samples with doses from 10 to 50 ng of mRNA (GFP). The results showed that N/P-1 and 10 are at doses 40 ng showed higher transfection of the mRNA which could be employed in future studies as a therapeutic dose.

The transfection of EPO with LNP-mRNA (EPO) in MG63 cells showed the ability of protein expression. The transient expression of the protein in the cytoplasm will be secreted by the cells into the media. The ELISA of the cell culture media with different loading of mRNA with fixing the N/P ratio to 5 showed a dose dependent expression of EPO protein (Figure 4A). However, the expression got saturated with the mRNA content of 62.50 ng which could be considered as a highest with the expression of 600 mU/mL. Figure 4B shows the viability of MG63 cells as 100% except for mRNA content 125 ng to 75% which is still higher than a 50% viability limit for a therapeutic.

## 4. Discussion

The application of MaGIC along with SMART has reduced the particle size significantly to approximately 50 nm with 100% encapsulation efficiency. This method is based on an additive manufacturing technology that allows for the designing and production of personalized particulate based therapeutic formulations with a precise control over the shape, size, and potentially the geometry.

The N/P ratio of 5 was set as an optimized formulation to balance the toxicity and therapeutic efficacy of EPO mRNA in MG63 cells *in vitro*. The amount of mRNA was further optimized to see the maximum therapeutic efficacy and low toxicity which yielded around 62 ng. The developed process would be able to develop robust and high mRNA encapsulation formulations at nanometer sizes that will have high penetrability and the ability to escape the ERS system in cancer. Hence the technology is the future for making robust doses of LNP-mRNA therapeutics for treating various diseases.

Erythropoietin (EPO) treats anemia in cancer patients undergoing chemotherapy. However, the same molecule also causes angiogenesis aggravating tumor growth. However American Society of Clinical Oncology (ASCO) recommended the usage of EPO in patients suffering with anemia with hemoglobin levels fall below 10 g/dL^37^. The administration of a single injection of mRNA EPO increased the serum EPO levels significantly in mice and lasted for more than a day^38^. A strong response in mice with a transient expression of gene of the is a promising development for further improvement. The naked gene administration always poses the risk of degradation by RNases and other factors.

EPO is a glycoprotein cytokine that acts as HGF and stimulates bone marrow erythropoiesis. It has been clinically prescribed for multiple conditions including anemia^39^. Several studies support the hypothesis that EPO promotes bone regeneration by inducing osteogenesis and angiogenesis. Trauma, infection surgical removal of tumor tissues and osteonecrosis resection lead to bone defects that can be treated using the novel bone regeneration approaches. Bone regeneration also greatly depends on osteogenesis as well as angiogenesis. Therefore, using EPO for bone remodeling purposes can be beneficial. Using mRNA for therapeutic purposes offers distinct advantages over traditional approaches involving proteins and recombinant proteins. mRNA-based therapies enable rapid and scalable production without the need for complex protein purification processes. This accelerates the development and manufacturing timelines. Additionally, mRNA provides a more direct route to intracellular protein production, facilitating precise control over the expression of therapeutic proteins. The flexibility of mRNA platforms allows for rapid adaptation to emerging pathogens and various diseases, showcasing their potential as a versatile and efficient tool in the realm of biotechnology and medicine. Using mRNA instead of proteins in various applications offers several other advantages such as reduced immunogenicity compared to protein therapies and simpler storage requirements compared to protein-based counterparts. Accordingly, using EPO mRNA in bone regeneration instead of EPO protein will be beneficial. It’s important to note that mRNA technologies also come with challenges, such as ensuring stability and delivery to target cells.

Nanoparticles play a pivotal role in mRNA delivery, enhancing their therapeutic potential. They protect mRNA from degradation, facilitate targeted delivery to specific cells or tissues, and improve cellular uptake. These properties are crucial for maximizing the efficacy of mRNA-based treatments, ranging from vaccines to personalized therapies in regenerative medicine. In this study, we fabricated lipid nanoparticles to encapsulate the negatively charged EPO mRNA and deliver it into the cells. Our results confirmed that mRNA was delivered into the cells successfully and the target protein, i.e. EPO, was expressed. Several methods have been developed for production of mRNA loaded LNPs. Examples of these methods include microfluidic devices, high-pressure or ultrasonication homogenizers and extruders. Although using these methods leads to a great degree of success, there is still a need to develop a method that can be easily integrated with other processing units to enhance the manufacturing of formulations and devices containing LNPs.

3D printing is an innovative approach that allows for the creation of customized scaffolds, promoting cell adhesion and growth. 3D-printed tissues hold immense potential for regenerative medicine, offering tailored solutions for repairing or replacing damaged organs and tissues. This technology is highly used in the field of regenerative medicine and tissue engineering as a key processing unit.

In this work, we disclose a novel device that can be integrated to 3D printing for simultaneous production of mRNA loaded scaffolds in one single step. Using this device, we demonstrated that we can successfully make uniform monodispersed LNPs by a pneumatic 3D bioprinter that is commonly used for fabrication of personalized scaffolds for tissue engineering. We utilized EPO mRNA as our model mRNA since it is demonstrated to be effective in osteogenesis and angiogenesis differentiation.

## 5. Conclusion

The SMART-MAGIC process has validated the advantages in preparing ∼50 nm LNP particles with mRNA loading of around 100%. The process also achieved around 100% loading efficiency. The fluorescence microscopy images showed the transfection of LNP-GFP (mRNA) at different amounts of mRNA. The LNP-EPO (mRNA) particles showed therapeutic efficacy against chemotoxicity *in vitro*. The SMART-MAGIC could be an alternative to the current nanoparticle fabrication process with both quality and custom manufacturing capabilities.

## 6. Acknowledgements

The authors thank UT Austin and College of Pharmacy for their core facilities. The authors also thank Dr. Rana Ghosh and his lab for providing fluorescence microscopy facilities.

## 8. Graphical Abstract

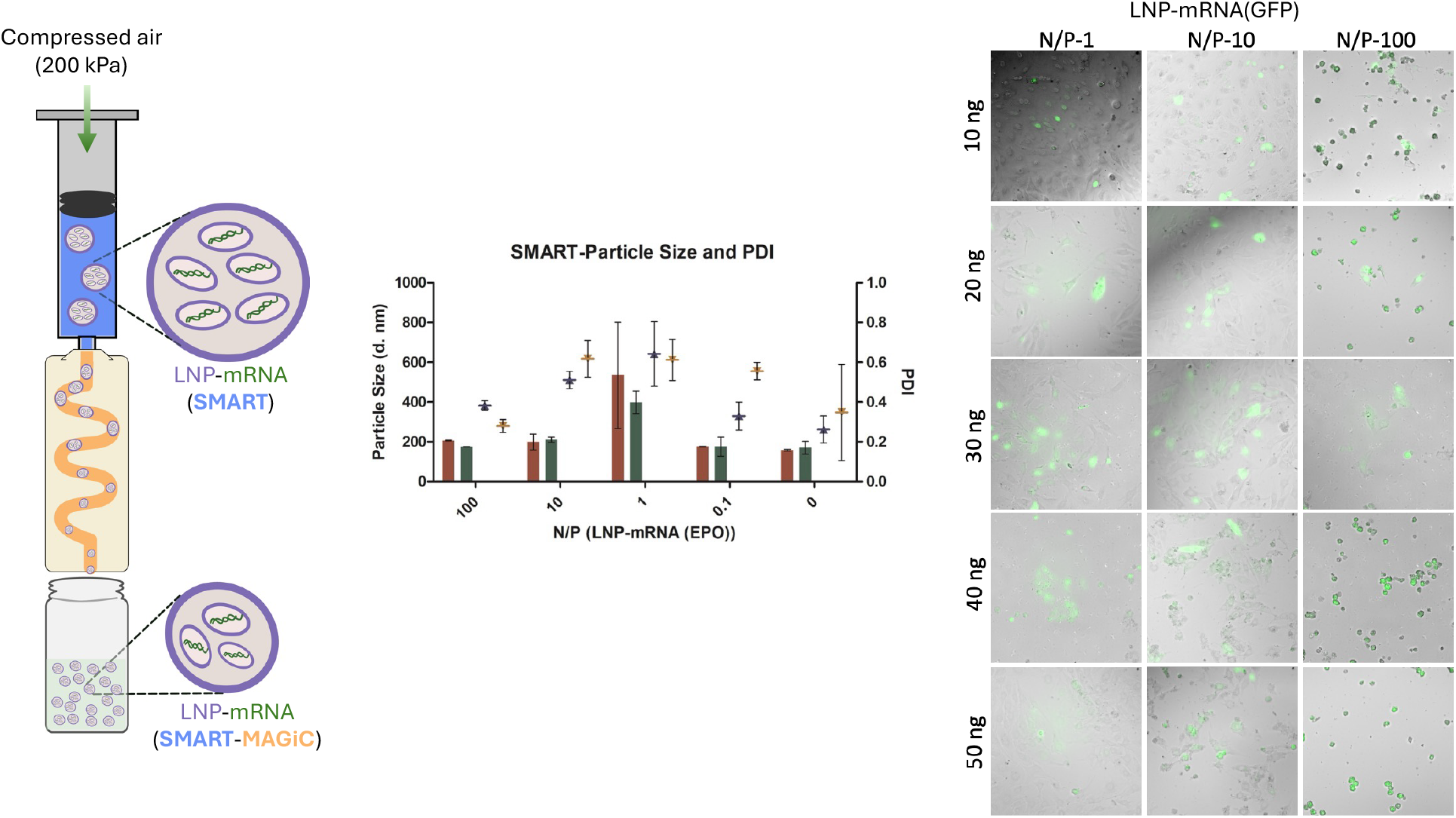

## 9. Supplementary Information

## Materials and Methods

### Sequence S1

#### GFP

L-7201 (capped with CleanCap AG, substituted with 5-methoxy-U) - https://www.trilinkbiotech.com/cleancap-egfp-mrna-5mou.html

1 AUGGUGAGCA AGGGCGAGGA GCUGUUCACC GGGGUGGUGC CCAUCCUGGU CGAGCUGGAC GGCGACGUAA ACGGCCACAA GUUCAGCGUG UCCGGCGAGG 101 GCGAGGGCGA UGCCACCUAC GGCAAGCUGA CCCUGAAGUU CAUCUGCACC ACCGGCAAGC UGCCCGUGCC CUGGCCCACC CUCGUGACCA CCCUGACCUA 201 CGGCGUGCAG UGCUUCAGCC GCUACCCCGA CCACAUGAAG CAGCACGACU UCUUCAAGUC CGCCAUGCCC GAAGGCUACG UCCAGGAGCG CACCAUCUUC 301 UUCAAGGACG ACGGCAACUA CAAGACCCGC GCCGAGGUGA AGUUCGAGGG CGACACCCUG GUGAACCGCA UCGAGCUGAA GGGCAUCGAC UUCAAGGAGG 401 ACGGCAACAU CCUGGGGCAC AAGCUGGAGU ACAACUACAA CAGCCACAAC GUCUAUAUCA UGGCCGACAA GCAGAAGAAC GGCAUCAAGG UGAACUUCAA 501 GAUCCGCCAC AACAUCGAGG ACGGCAGCGU GCAGCUCGCC GACCACUACC AGCAGAACAC CCCCAUCGGC GACGGCCCCG UGCUGCUGCC CGACAACCAC 601 UACCUGAGCA CCCAGUCCGC CCUGAGCAAA GACCCCAACG AGAAGCGCGA UCACAUGGUC CUGCUGGAGU UCGUGACCGC CGCCGGGAUC ACUCUCGGCA

701 UGGACGAGCU GUACAAGUAA

### Sequence S2

#### EPO

L-7209 (capped with CleanCap AG, substituted with 5-methoxy-U) - https://www.trilinkbiotech.com/cleancap-epo-mrna-5mou.html 1 AUGGGCGUGC ACGAGUGCCC CGCCUGGCUG UGGCUGCUGC UGAGCCUGCU GAGCCUGCCC CUGGGCCUGC CCGUGCUGGG CGCCCCCCCC CGGCUGAUCU 101 GCGACAGCCG GGUGCUGGAG CGGUACCUGC UGGAGGCCAA GGAGGCCGAG AACAUCACCA CCGGCUGCGC CGAGCACUGC AGCCUGAACG AGAACAUCAC 201 CGUGCCCGAC ACCAAGGUGA ACUUCUACGC CUGGAAGCGG AUGGAGGUGG GCCAGCAGGC CGUGGAGGUG UGGCAGGGCC UGGCCCUGCU GAGCGAGGCC 301 GUGCUGCGGG GCCAGGCCCU GCUGGUGAAC AGCAGCCAGC CCUGGGAGCC CCUGCAGCUG CACGUGGACA AGGCCGUGAG CGGCCUGCGG AGCCUGACCA 401 CCCUGCUGCG GGCCCUGGGC GCCCAGAAGG AGGCCAUCAG CCCCCCCGAC GCCGCCAGCG CCGCCCCCCU GCGGACCAUC ACCGCCGACA CCUUCCGGAA 501 GCUGUUCCGG GUGUACAGCA ACUUCCUGCG GGGCAAGCUG AAGCUGUACA CCGGCGAGGC CUGCCGGACC GGCGACCGGU GA

